# Evaluation of CD8 T cell killing models with computer simulations of 2-photon imaging experiments

**DOI:** 10.1101/830505

**Authors:** Ananya Rastogi, Philippe Robert, Stephan Halle, Michael Meyer-Hermann

## Abstract

*In vivo* imaging of cytotoxic T lymphocyte (CTL) killing activity revealed that infected cells have a higher observed probability of dying after multiple contacts with CTLs, suggesting memory effect in CTLs or infected cells. We developed a three-dimensional agent-based model of CTL killing activity to discriminate different hypotheses about how infected cells get killed based on quantitative 2-photon *in vivo* observations. We compared a constant CTL killing probability with mechanisms of signal integration in CTL or infected cells. The most likely scenario implied increased susceptibility of infected cells with increasing number of CTL contacts where the total number of contacts was a critical factor as opposed to signal integration over many contacts. However, when allowing *in silico* T cells to interact with apoptotic target cells (zombie contacts), a contact history independent killing mechanism was also in agreement with the experimental datasets. We showed that contacts that take place between CTLs and dying infected cells impact the observed killing dynamics because even in absence of modulation of cell properties, we saw an increase of the observed probability of killing infected cells with more interactions. The duration taken by an infected cell to die and the per capita killing rate (PCKR) of CTLs, parameters hard to measure directly, were determined from the model and turned out predictive to distinguish the different CTL killing models in future experiments. The comparison of observed datasets to simulation results, revealed limitations in interpreting 2-photon data, and provided prediction for additional measurements to distinguish CTL killing models.

**Highlights:**

- Killing of infected cells by cytotoxic T cells typically involves more than a single contact.
- Cytotoxic T cells or infected cells integrate signals from multiple interactions.
- T cell contacts with dying infected cells have a major impact on *in vivo* data interpretation.

**Significance Statement:** Despite having a clear understanding of cytotoxic T lymphocyte (CTL) mediated cytotoxicity mechanisms, the quantitative dynamics remain unexplored at a cellular level. We developed an agent-based model to compare different hypotheses for mechanisms of CTL mediated cytotoxicity that could lead to an increase in observed probability of killing infected cells at higher interactions with CTLs as seen *in vivo*. We showed that this behaviour can be explained by modulation of properties by infected cells or CTLs with increasing number of contacts. For the modulation, we compared two modes of signal integration and showed that time is not a relevant parameter in signal integration. We also studied the impact of contacts between CTLs and apoptotic infected cells on observed killing properties.

## Introduction

Protection from intracellular pathogens like viruses requires the detection and clearance of intracellularly infected cells. One of the most prominent cell-mediated protective immune effector mechanisms is the cytotoxic T lymphocyte (CTL) response. CTLs migrate within lymphoid tissues and can travel between infected organs and search for infected cells presenting foreign antigens. The motility properties, the capacity to find infected cells, and ultimately the decision to kill infected cells with minimal damage to healthy cells are critical factors for an optimal containment of infection.

Upon target cell encounter, CTLs can initiate contacts with infected cells that lead to the formation of an immunological synapse through which CTLs deliver perforin and granzymes to kill the infected cells ([1], [2]). Various mechanisms can be used by CTLs to kill infected cells, which include secretion of cytotoxic granules, surface-receptor mediated signalling, and secretion of cytokine [3]. Many of these protective functions of CTLs have been explored and are well understood [4]. However, until recently, most of these studies relied on *in vitro* CTL assays and on cell culture techniques. *In vitro* studies of killing dynamics at a cellular level have revealed that infected cells form simultaneous contacts with multiple CTLs [5].

To better quantify the results observed in experimental systems, analyses have been performed to calculate the per capita killing rate (PCKR) of CTLs and the death rate of infected cells. These include differential equation–based approaches with various spatial compartments described by their concentrations ([6], [7], [8], [9]). In a series of studies consisting of a simulated environment made to resemble a portion of the lymph node, Gadhamsetty et al. have shown that a double saturation function is the best fit for the killing rate ([10], [11]). This was done with a 2D cellular Potts model and was used to explore killing regimes such as monogamous killing, where one CTL can kill just one infected cell at a time; simultaneous killing, where one CTL can kill multiple infected cells at the same time; joint killing, where multiple CTLs kill a single infected cell; and mixed killing, where multiple CTLs can kill infected cells simultaneously.

To understand the dynamics of how CTLs interact and kill infected cells in a three-dimensional (3D) tissue *in vivo*, Graw et al. [12] developed a 3D model to look at CTL mediated killing through the lens of CTL motility. Indeed, the dimensionality of Potts models affects the efficiency of CTL mediated killing of target cells [13]. These models use an empirical probabilistic mechanism. The killing mechanisms at an individual cellular level remained unresolved.

Previously, we reported *in vivo* CTL-mediated killing kinetics analysed by 2-photon microscopy *in vivo*. We tracked CTLs interacting with virus-infected cells *in vivo* [14] inside lymph nodes with fluorescent reporter viruses that allow direct observation of the infected target cell over time. These studies relied on morphological disruption of the target cell as evidence for irreversible target cell death and showed that one CTL contact event typically did not suffice to achieve killing. Instead, the experimental data showed that target cells with more than one interaction with CTLs during the observation period were more frequently disrupted than target cells with one interaction. The observed probability of killing infected cells also showed an increase with increasing number of interactions with CTLs. This observation challenged the conventional view of CTL immunity, that only one contact suffices, and further suggests complex mechanisms by which CTLs adapt their killing efficiency or infected cells become more susceptible to death.

Here, we developed an agent-based model that reproduces the *in vivo* movement and interactions of CTLs with infected cells in 3D (see methods) and tested various hypotheses about which killing mechanisms at the cellular level can explain the quantitative killing dynamics observed in [14]. For each of the hypotheses, we also explored the impact of contacts between already apoptotic infected cells and CTLs, which we termed as “zombie contacts”.

The results from the model showed that retention of information about prior contacts, by either CTLs or infected cells, results in an increase of the observed probability of killing infected cells with more interactions. Strikingly, the presence of zombie contacts challenged the interpretation of CTL killing activity, because even when infected cells and CTLs do not modulate their properties, we saw an increase of the observed probability of killing infected cells with more interactions. With this model, we estimated the time taken by a cell to disappear once the decision for death has been taken and the PCKR. The model unravelled that different types of mechanisms exhibit different PCKR profiles suggesting that PCKR measurements could further pin down the mechanism at work.

## Results

### Agent-based model of T cell-mediated killing

In order to dissect the CTL-mediated killing activity observed *in vivo* in [14], we developed an agent-based model that reproduces the movement and interactions of CTLs with non-motile virus-infected cells in a three-dimensional context similar to the *in vivo* imaging setup [14] (Figure 1a) (see Methods). We modelled CTL migration to mimic T cell behaviour as observed by 2-photon microscopy (Figure 1b) and once the CTLs and the virus-infected cells are at a threshold distance from each other, an interaction is initiated (Figure 1c). The CTLs move in a direction and with a speed that we randomly picked from measured distributions (Figure 1d, e). After each persistent time, a new direction and speed are sampled from those distributions (see Methods). We focused on the case of CTLs reacting to antigen-expressing virus-infected cells with a relatively low average CTL speed [14]. The duration of an interaction is directly taken from experimental data (Figure 1f). The resulting model quantitatively reproduces CTL migration in the case of antigen recognition by peptide-MHC class I presentation on virus-infected cells.

**Figure 1:**
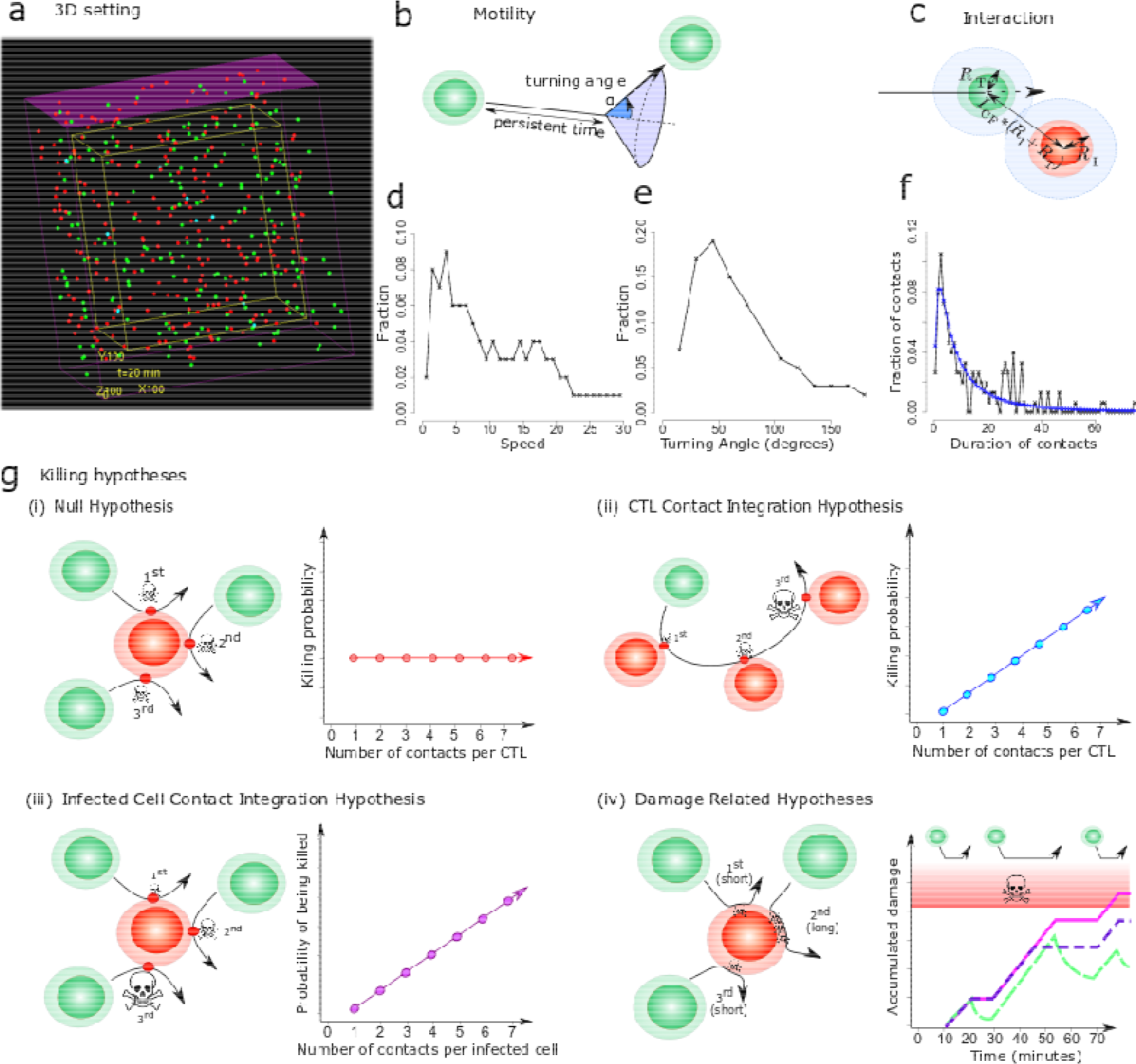
Design of the agent-based model for spatial CTL killing activity. (a) 3 dimensional view of the agent-based model, (b) CTLs in the model move in a straight direction until a persistence time is reached, then a new direction is randomly set according to an experimentally measured turning angle distribution, (c) a CTL interacts with an infected cell when they are in close proximity to each other, (d) speed distribution and, (e) turning angle distribution of CTLs obtained from experiments and used in the model, (f) distribution of contact durations between infected cells and CTLs (black-experimental data, blue- log normal plot fitted to experimental data and used in the model), (g) different killing hypothesis: (i) Null hypothesis: Infected cells and CTLs do not retain memory of prior contacts. Each contact is associated with a constant probability of death. The associated plot shows the behaviour of CTL killing properties with more interactions with infected cells; (ii) CTL contact integration: The CTLs retain memory of contacts and with each subsequent contact, CTLs become more lethal. The associated plot shows the behaviour of CTL killing properties with more interactions with infected cells; (iii) Infected cell contact integration: The infected cells retain memory of CTL contacts. The associated plot shows the behaviour of infected cell susceptibility to death with more interactions with CTLs; (iv) Damage hypotheses: The CTLs induce damage to the infected cells. The associated plot shows the behaviour of damage of infected cells with more interactions with CTLs for different hypotheses: damage (pink), damage and repair (green) and saturated damage (purple). The red area represents the damage greater than the threshold damage after which an infected cell dies.

Using this basic model setup, we compared different hypotheses regarding possible CTL-mediated killing mechanisms (Figure 1g). The (i) *Null hypothesis*: defined as a scenario with a contact history-independent CTL-mediated target cell killing, i.e. CTLs kill with equal probability at each CTL-target cell contact. The considered alternative hypotheses are: (ii) *CTL contact integration*: modulated killing capacity of CTLs with more CTL-target cell interactions; (iii) *infected cell contact integration:* increased susceptibility for death at higher number of contacts with CTLs; and (iv) *damage:* damage accumulation of infected cells with each contact with CTLs, possibly combined with repair of the cell, which is the *damage and repair hypothesis*, or with a limited damage per contact due to T cell exhaustion, which is *saturated damage*. The datasets used to discriminate between killing hypotheses are taken from [14] (Figure S1).

### A contact history independent killing mechanism is in contradiction to 2-photon experiments

First, we investigated whether the observed properties of CTL-mediated killing seen in quantitative 2-photon imaging (see Supplement, Figure S1) could be explained by a contact history independent killing mechanism (Null hypothesis). Although most of the model's parameters are directly known from experimental measurements, the probability of a CTL to kill an infected cell and the time between activation of apoptosis and visual cell dissolution remained to be estimated. The parameter values best reflecting the experimental data were identified. Only a restricted range of parameters was consistent with the data (see Methods and Supplement, Figure S2a).

As expected in this model scenario (see Figure 2a-h, green lines), the observed probability of killing infected cells (see Methods) remained constant with increasing number of interactions (Figure 2a, green line), which is in contradiction to the experimental results (black lines). Thus, testing the Null hypothesis serves as a sanity check that the model works properly and that the observed *in silico* probability of killing infected cells with more interactions reflects the cellular CTL-mediated killing mechanism in this context. Further, it shows that a simple contact history independent killing probability cannot explain the *in vivo* datasets of Halle et al. with this model setting [14].

**Figure 2:**
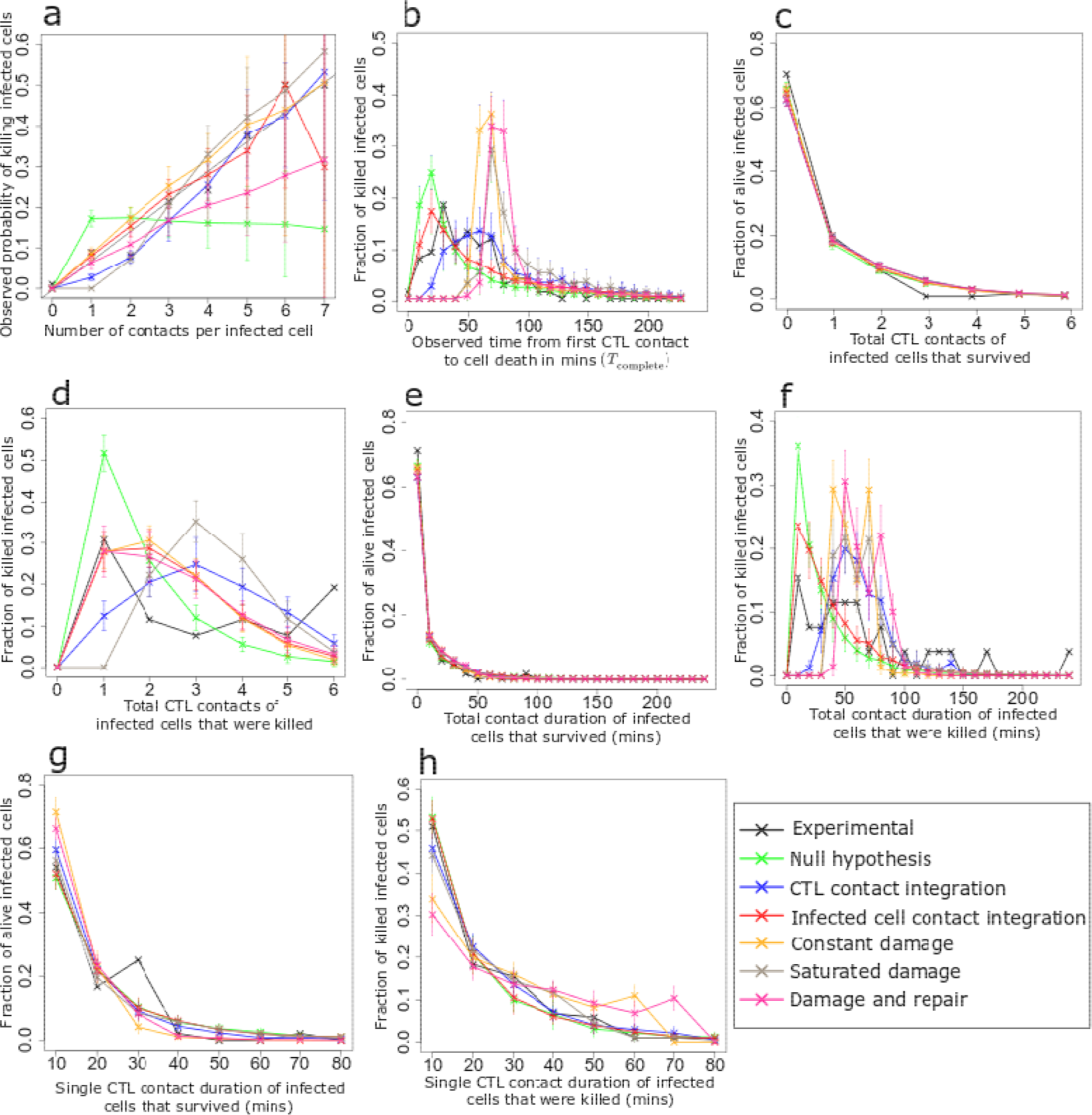
Analysis of *in silico* killing simulations in the absence of zombie contacts. Simulation results are compared with experimental measurements for different killing hypotheses in the absence of zombie contacts, using the best identified parameters for each hypothesis (Figure S2). (a) Observed probability of killing infected cells in dependence on the number of interactions with CTLs, (b) distribution of observed times between first contact to a CTL and actual cell death for all killed infected cells, (c) distribution of the number of contacts with CTLs for all infected cells that survived during the observation period and, (d) were killed during the observation period, (e-h) distribution of total (e, f) and single (g, h) contact durations with CTLs for infected cells that survived during the observation period (e, g) and were killed during the observation period (f, h). Error bars represent SD from 30 simulations.

### Contact integration mechanisms on infected cells or CTLs are compatible with 2-photon experiments

In order to explain the increase in observed probability of killing infected cells with each interaction with CTLs (Figure 2a, black line), we hypothesized that this could reflect a modulation of CTL killing capacity each time they encounter an infected cell (Figure 1g(ii)). In this hypothesis, CTLs kill with a linearly increasing probability proportional to the number of previous target cell contacts of this CTL. The cost (quality) of simulations with different parameter values is shown, and an optimal parameter set could be identified (see Supplement, Figure S2b).

For this optimal parameter set, the observed probability of killing infected cells in dependence on the number of interactions with CTLs and the corresponding analyses for the other experimental datasets (Figure 2a-h, blue lines) showed a good agreement with the experimental results. Thus, the hypothesis that CTLs modulate their killing capacity is compatible with the experimental observations.

Next, we explored whether infected cells might become more sensitive to cytolysis with increasing numbers of CTL contacts (infected cell integration hypothesis, Figure 1g(iii)). The death susceptibility of infected target cells was modelled as a linearly increasing probability of death that is proportional to the number of prior CTL visits (see methods and Supplement, Figure S2c). The simulation with the best parameter set (Figure 2a-h) is also in agreement with the experimental data suggesting that infected cells may get more susceptible to cell death with increasing number of CTL contacts.

### Damage accumulation of infected cells at a linear rate is not compatible with 2-photon experiments

Having established that infected cells retaining memory of prior cell contacts is a possible process, we wanted to elucidate the mechanisms by which infected cells could retain memory of previous CTL contacts. We hypothesized that infected cells get damaged by interacting CTLs and once the damage of an infected cell reaches a threshold value of 1 (100%), the cell dies. The three damage-based hypotheses that were considered were constant damage, saturated damage, and damage and repair (Figure 1g(iv)). While all other hypotheses have one parameter to describe the killing dynamics of infected cells by CTLs, saturated damage hypothesis and damage and repair hypothesis have two parameters.

The constant damage hypothesis assumes a constant rate of damage during each interaction. In the saturated damage hypothesis, the damage process by the CTL stops after the interaction exceeds a threshold time (*T*_max_). This hypothesis was proposed because CTLs have a storage of cytolytic granules and are possibly not able to sustain a damage process for very long interactions, reaching up to 40 minutes in the *in vivo* dataset. Therefore, a possible biological factor that affects the killing observations could be T cell exhaustion during single contacts. We assumed that the time lapse in between two contacts is long enough to allow the CTLs to recuperate and damage the next contacted infected cells again. In the damage and repair hypothesis, there is no T cell exhaustion. The CTLs damage the infected cells throughout the duration of the interaction but the infected cells also repair themselves between different CTL visits.

The increase in observed probability of killing infected cells with increasing number of interactions with CTLs was recapitulated for the constant and the saturated damage hypothesis (Figure 2a, orange and brown line) with the best parameter set in Figure S2d and e. However, the simulated observed time between the first contact to a CTL and the actual cell death (*T*_elimination_) did not coincide with the experimental distribution (Figure 2b, orange and brown line).

The best damage rate was approximately *d* = 0.03 per minute for the constant and the saturated damage hypotheses. In order to achieve a damage level of 1, the total contact time required would be in the range of 30 minutes (see Methods) which is the minimum value for *T*_elimination_. This is a lower bound because we assumed that the time between the fate decision for death and the actual dissolution of the cell is negligible. For the saturated damage hypothesis, the value would exceed these 30 minutes (see Methods). For the damage and repair hypothesis, the optimal value for the damage rate was 0.03 damage per minute and the value for the repair rate was 0.009 per minute (Figure S2f). The duration of one contact sufficient to induce a total damage of 1 was approximately 40 minutes (see Methods, equation (5)). For an infected cell that died, *T*_elimination_ will also include the time that lapsed between consecutive contacts. As shown by the calculations above, for all of the damage-based hypothesis, no infected cell can have a *T*_elimination_ value less than 30 minutes which is in disagreement with the experimental data (Figure 2b). Thus, we conclude that none of the tested damage-based hypotheses are compatible with the experimental datasets.

### Constant killing probability with the assumption of zombie contacts is compatible with 2-photon experiments

Virus-specific CTLs may interact with target cells even during the process of target cell death. Thus, a target might receive multiple CTL contact events, even though the fate decision to die has already been taken (“zombie contacts”). However, we have observed that once a target cell is disrupted, the remaining small remnants of the target cells are contacted by CTLs only sometimes. We argued that most of these remnants are taken up by local dendritic cells and macrophages, thus preventing direct access of the CTLs to the membrane of the remnants.

It is unknown whether CTL interactions with target cells in the process of cytolysis might affect the analysis of CTL killing mechanisms and estimates of CTL killing efficiency. To address this question, we next tested whether the allowance of such zombie contacts impacts the observed probability of killing infected cells with increasing number of interactions with CTLs.

Simulations with the Null hypothesis with zombie contacts were performed with different killing probabilities and times for infected cell death. An optimal parameter set could be identified (see Supplement, Figure S3a). The simulations (Figure 3a-h, green lines) show that, although CTLs mechanistically kill with equal probability at each contact in the model, in the presence of zombie contacts the observed probability of killing infected cells exhibits an increase with increasing number of interactions with CTLs (Figure 3a, green line).

**Figure 3:**
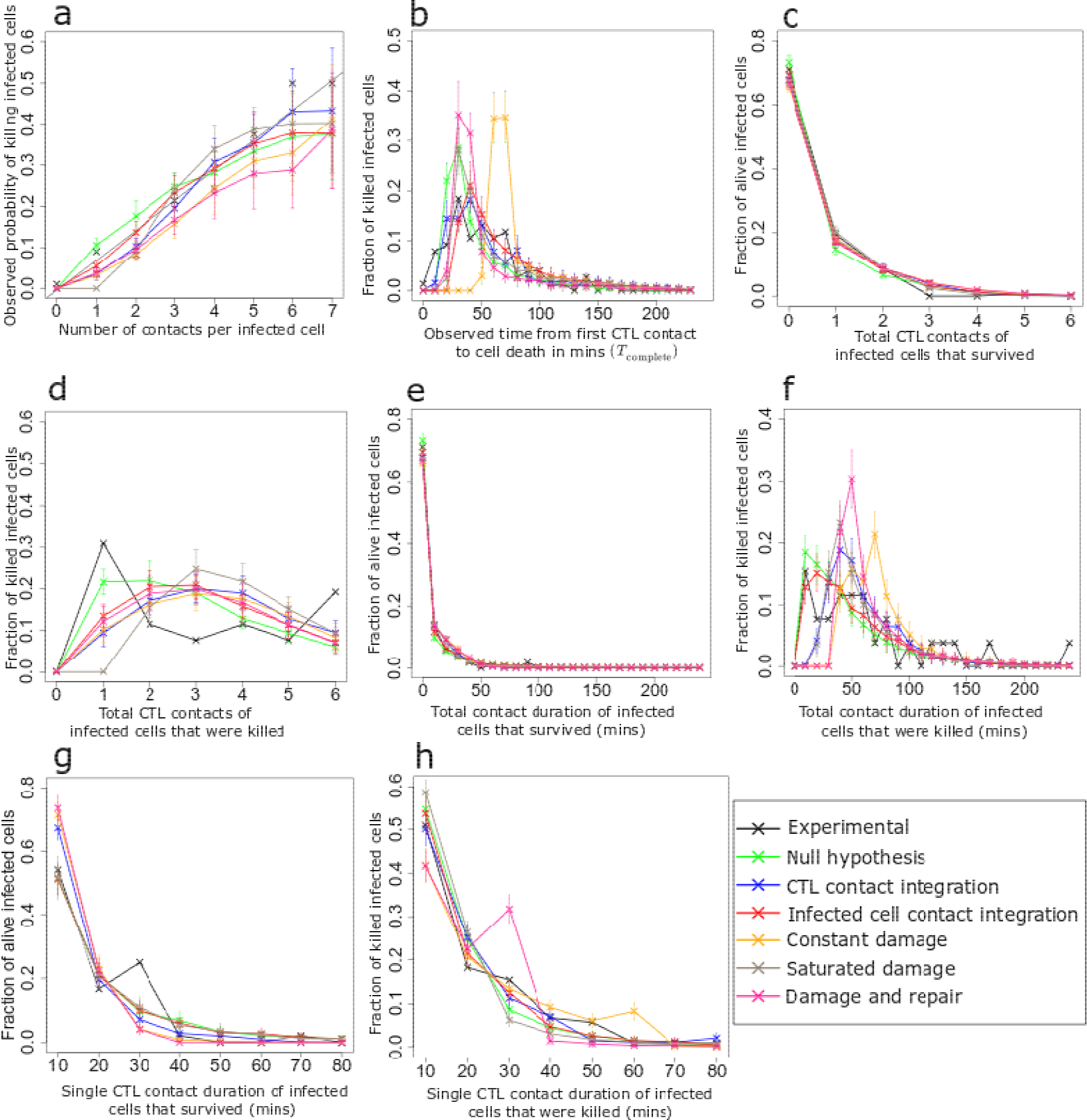
Analysis of *in silico* killing simulations in the presence of zombie contacts. Simulation results compared with experimental measurements for different killing hypotheses in the presence of zombie contacts, using the best identified parameters for each hypothesis (Figure S3). Curves are depicted similar to Figure 2. Error bars represent SD from 30 simulations.

Due to zombie contacts, when following CTL killing activity *in vivo*, the time during which cells are already dying is counted in the analysis of the number of contacts. This leads to an over-estimation of the number of contacts compared to the real number of contacts that lead to the decision for cell death. Therefore, we concluded that the increase in observed probability of killing infected cells with each CTL interaction can be directly influenced by zombie contacts, even when the CTLs and infected cells do not adapt their properties. Interestingly, the Null hypothesis in presence of zombie contacts was compatible with all other datasets (Figure 3b-h, green lines). In addition, after the 6^th^ contact, the simulations showed a saturation in the observed probability of killing infected cells while the experimental dataset continues increasing. Eventually, a longer observation time would increase the total number of observed contacts per target cells and allow further supporting or discarding the Null hypothesis as possible mechanism.

### Zombie contacts have a major impact on the interpretation of 2-photon experiments and on model selection

Considering the observed relevance of zombie contacts in the Null hypothesis, we revisited all other hypotheses to test whether addition of zombie contacts to our model impacts model performance in other scenarios as well (Figure S3b-f). CTL contact integration and infected cell contact integration gave results in agreement with the experimental results; but none of the damage-based hypotheses gave rise to plots that were in agreement with the experimental results. This implies that apart from the Null hypothesis, the ranking of the model hypotheses remained the same irrespective of the presence or absence of zombie contacts. However, all hypotheses showed a lower cost and a lower Akaike information criterion (AIC) value in presence of zombie contacts (Figure 4a, Table 1, 2).

**Figure 4:**
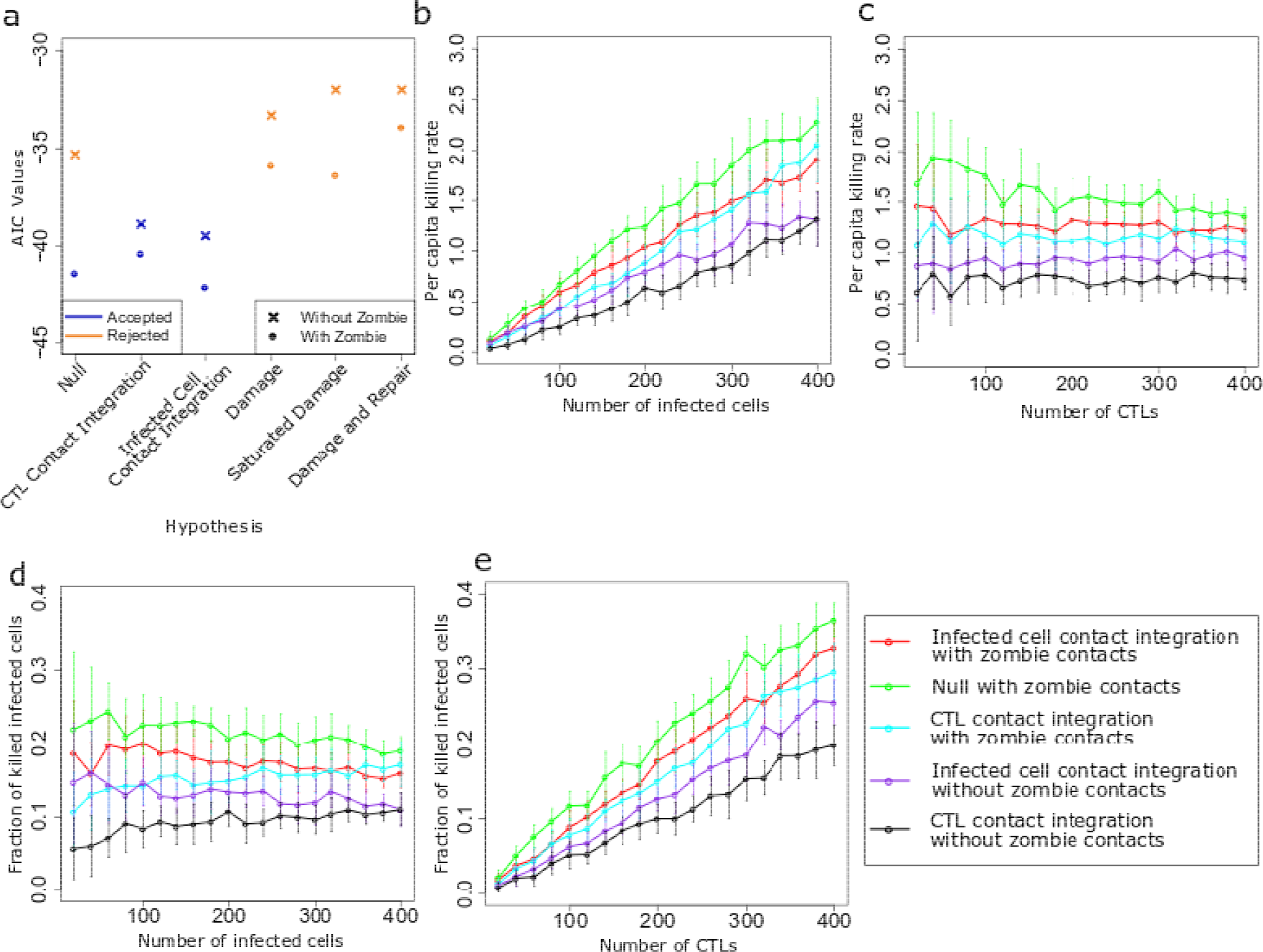
Model selection and parameter prediction. (a) Comparison of AIC values for all hypotheses corresponding to their respective least cost. The data points in blue show the hypotheses that gave a good agreement with experimental results. The data points in orange show the hypotheses that were rejected. (b, c) PCKR values for variable number of infected cells (b) and of CTLs (c), (d, e) Fraction of killed cells for variable number of infected cells (d) and of CTLs (e). Error bars represent SD from 30 simulations.

**Table 1:** Least cost for all hypotheses in absence of zombie contacts in ascending order of AIC

| Hypothesis | Killing parameter | $T_{\text{death}}$ | Least cost | AIC |
| --- | --- | --- | --- | --- |
| Infected cell contact integration | 0.09 | 15 | $4.35 \times 10^{-3}$ | -39.5 |
| CTL contact integration | 0.08 | 10 | $4.70 \times 10^{-3}$ | -38.9 |
| Null hypothesis | 0.2 | 15 | $7.35 \times 10^{-3}$ | -35.3 |
| Constant damage | 0.03 | 35 | $9.37 \times 10^{-3}$ | -33.3 |
| Damage and repair | $d = 0.03, r = 0.009$ | 40 | $8.60 \times 10^{-3}$ | -32.0 |
| Saturated damage | $d = 0.03, T_{\text{max}} = 20$ | 35 | $8.64 \times 10^{-3}$ | -32.0 |

**Table 2:**
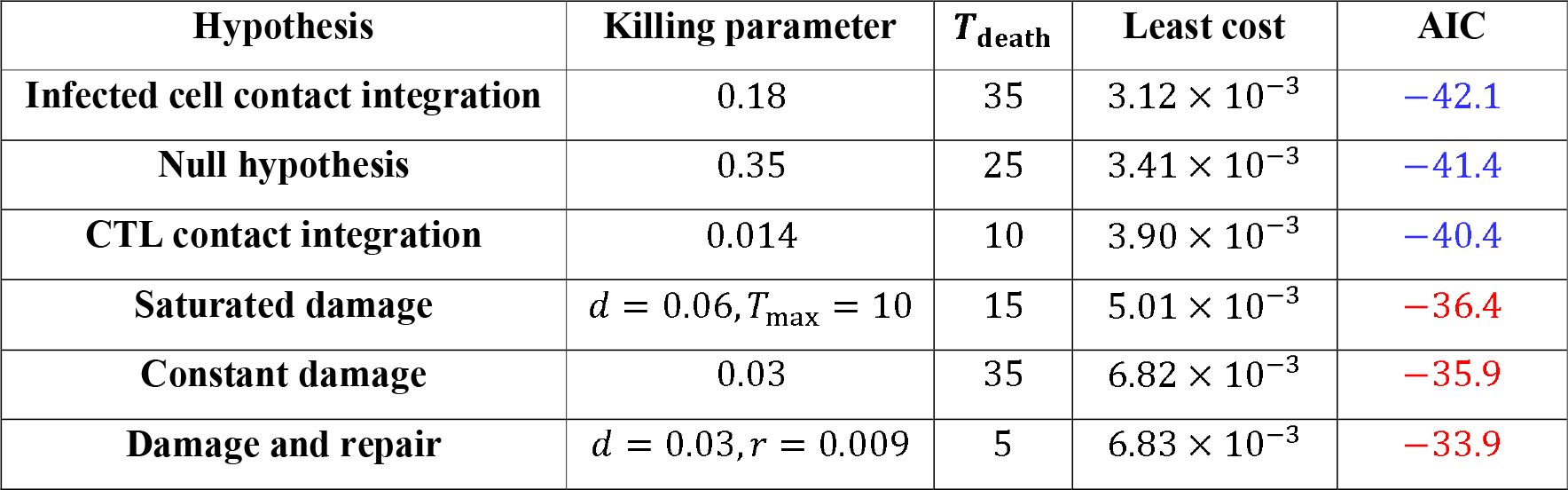
Least cost for all hypotheses in presence of zombie contacts in ascending order of AIC

### The per-capita killing rate and the fraction of killed cells can discriminate hypotheses

By selecting the optimal parameter and plotting the readouts obtained from the experiments, various hypotheses were discarded. The five hypotheses that give a good agreement with all datasets (Figure 4a) in ascending order of cost are: (i) Infected cell contact integration in presence of zombie contacts, (ii) Null hypothesis in presence of zombie contacts, (iii) CTL contact integration hypothesis in presence of zombie contacts, (iv) infected cell contact integration in absence of zombie contacts, and (v) CTL contact integration hypothesis in absence of zombie contacts.

We sought to predict properties of the different hypotheses that could further help to discriminate them or to design new predictive experiments. One property is the observed PCKR of CTLs, as a measure of 3D population killing efficiency. The PCKR is defined as the number of infected cells killed per CTL in 24 hours. Another important measure is the fraction of killed infected cells defined as the ratio of killed infected cells to the total number of infected cells in the system.

To understand how the number of infected cells affects the PCKR values, the number of CTLs was kept constant at 200 while the infected cell number was varied from 20 to 400 (Figure 4b). The other parameter values were kept fixed at the optimal parameter set obtained from the respective cost heatmaps of each hypothesis. The increase of PCKR values was highest for the Null hypothesis but for all five hypotheses, the values of PCKR increased with increasing number of infected cells.

Next, to elucidate the effect of the number of CTLs on the PCKR, the number of infected cells was kept constant at 250 while the CTL numbers were varied from 20 to 400 (Figure 4c). Surprisingly, even when the ratio of infected cells to CTLs is high, the values of PCKR remained constant for all hypotheses except the Null hypothesis for which we saw a small drop in values of PCKR with increasing number of CTLs.

In the same spirit, we investigated the fraction of infected cells that are killed by CTLs for different initial numbers of infected cells. A surprising result is that even for a low number of infected cells, CTLs fail in killing all infected cells within the simulated 4-hours (Figure 4d). In addition, the fraction of killed cells increased with the initial number of infected cells for the CTL contact integration hypothesis, irrespective of zombie contacts, while it reduced for all other hypotheses.

In contrast to the results obtained for variable number of infected cells, the fraction of killed infected cells increased with number of CTLs for all hypotheses (Figure 4e). At higher numbers of CTLs, a much larger fraction of infected cells is killed. Taken together, these results suggest that by keeping all other factors of a system constant and varying either the number of infected cells or the number of CTLs, we can differentiate between different killing hypotheses by analysis of PCKR and fraction of killed cells.

## Discussion

Following up on in vivo 2-photon microscopy-based imaging of CTL-mediated immunity [14], we developed a 3-dimensional agent-based model to visualize and quantify the dynamics of virus-infected cells and CTLs. Different hypotheses on how CTLs kill infected cells were compared to observed CTL killing dynamics. Cells other than infected cells that might be able to interact with CTLs by processing and presenting extracellular antigens with MHC class I molecules to them were not included. Already with this restricted model complexity, a combination of measured datasets was required to discriminate different hypotheses.

To discriminate between the hypotheses, we evaluated a cost between simulations and all combined datasets. While the experimental data for observed probability of killing infected cells included data for up to 14 contacts with CTLs, the experimental data points beyond the 6^th^ contact only represent a small number of cells. Due to this, only the 6 first interactions are included in the cost calculation. Thus, experiments with more data points at higher CTL contact numbers could help in model selection.

For all hypotheses, a better fit with experimental data was found in the presence of zombie contacts, suggesting that there is a substantial time between the decision of death and actual disappearance of the cell, and that it is critical to include this parameter. Importantly, there is no reliable *in vivo* reporter system for this hypothetical mechanism available. Notably, caspase activity reporter systems might be useful [15], but it is typically not clear at what stage of caspase activity the pro-cell death pathway passes an “irreversibility threshold” in the context of an ongoing CTL attack. Thus, better *in vivo* sensors of target cell viability will be helpful to better define the “point of no return” of CTL-mediated target cell killing.

We showed that retention of information about prior contacts, by either CTLs or infected cells, is compatible with all the observed datasets. Surprisingly, in the presence of contacts between dying infected cells and CTLs (zombie contacts) a contact history independent killing mode (Null hypothesis) gave rise to an increase in the observed probability of killing with each contact of the infected cell with a CTL. This raises the question if in the experimental system, the killing is really getting more efficient or is an artefact of zombie contacts. Thus, the hypothetical possibility of zombie contacts introduces another uncertainty into the analysis of live-imaging data. To further enhance our understanding of this system, it is imperative to be able to distinguish zombie contacts from other contacts in an experimental setup. In the case that zombie contacts actually happen, the present simulations might be used to reconstruct the real killing efficiency given the measured data.

In the simulations, we sampled the duration of contacts from just one distribution. This implicitly assumes that CTL contacts with zombie and alive cells are of equal duration. The possibility of two distinct distribution of duration of contacts could be tested through aforementioned sensors to detect apoptotic phases of infected cells. Inclusion of different distributions of contact durations may further progress our understanding of the system.

We established that a positive modulation of CTL killing with each contact with infected cells is a viable hypothesis. However, CTLs in the context of chronic infections or cancer can get exhausted ([16], [17]). Exhaustion is usually defined as a reduced protective response in the face of ongoing antigenic activation. E.g. during virus infections exhausted CTLs lose activities like direct killing and cytokine production ([18], [19]). These mechanisms are in general viewed as negative modulation of the killing capacity of CTLs [20]. Thus, on the single cell level, a CTL that interacted with a sequence of multiple target cells would rapidly become less cytotoxic, being able to kill fewer-and-fewer target cells. Interestingly, none of the parameter values tested here supported such a decreased killing efficacy of infected cells per contact (data not shown). This suggests that, at the time point studied and within the duration of analysis and in this experimental setting, exhaustion at the time-scale of multiple contacts is not dominant. On the contrary, the simulation suggests that CTLs get more lethal with increasing numbers of contact. The CTL contact integration hypothesis naturally gives rise to a diverse population of CTLs that show heterogeneity during killing. This is in agreement with studies showing that CTLs have dissimilar killing properties that contribute to a robust T cell response ([21], [22]).

Signal integration from multiple CTLs on the side of infected cells was also found in agreement with the experimental data. We, thus, proposed damage-based hypotheses as possible mechanisms by which the retention of contacts by infected cells is implemented. Interestingly, the hypotheses of damage, saturated damage and concomitant damage and repair are not consistent, because a too low damage rate was imposed by the data (Figure S1a) leading to a contradiction of the experimental and simulated time between the first contact to a CTL and actual cell death, *T*_elimination_ (Figure S1b). While the infected cell contact integration hypothesis describes a system where the number of CTL contacts determines cell death, the damage-based hypotheses also resolve the duration of the interactions, i.e. a longer interaction leads to a greater damage ([23], [24], [25]). The simulation results suggest that the number of contacts with CTLs rather than the time integration of signals is critical for the target cell fate decision.

The 2-photon imaging datasets (Figure S1), despite encompassing multiple levels of information on duration of contacts and tracking of cell contacts, was not sufficient to identify whether the CTL-mediated killing rate is constant or integrated by the CTL or infected cell. However, we here showed that more indicators can discriminate these remaining hypotheses. A first indicator is the killing efficiency of CTLs, as defined by PCKR of CTLs [26]. The PCKR values obtained for the final hypotheses are different for varying number of initial CTLs and infected cells. This suggests that a careful measurement of PCKR values for different cell numbers is suitable to discriminate them, and would need to be performed with comparable numbers of CTLs and infected cells (Figure 4b-c).

A recent study [27] reported a higher PCKR value at low CTL to infected cell ratio. In the simulations, we observe a similar behaviour when we vary the number of infected cells while keeping the number of CTLs constant (Figure 4b). But a higher PCKR value at low CTL to infected cell ratio was not in agreement with the results obtained when keeping the number of infected cells constant and varying the number of CTLs. Thus, our results suggest that instead of the ratio of CTLs to infected cells, the number of infected cells impact the PCKR values.

At very low numbers of simulated infected cells, the CTLs still fail to kill all infected cells which contradicts the rapid CTL mediated killing seen in other studies [28]. This could be a consequence of the assumed random cell movement and raises the question whether directed cell movement where CTLs actively migrate towards infected cells is important to ensure successful elimination of more infected cells.

In [14], the PCKR values were found to range from 2 to 16. It is interesting that with our model, the hypothesis that gives the values most consistent with this range is the Null hypothesis in the presence of zombie contacts. While this information is not enough yet to reject the other hypotheses, it gives an interesting insight into the behaviour of CTL mediated killing. The significance of cytotoxicity mediated by CTLs and the impact of the number of cells on PCKR and fraction of killed infected cells are especially important due to the role of T cells in viral infections such as HIV, viral pneumonia and other diseases such as cancer ([29], [30]).

The fraction of total killed infected cells shows a linear increase with increasing number of CTLs, implying that CTL mediated killing could follow mass action killing kinetics. Previous studies have shown that T cells show mass action killing kinetics in the spleen [31] and our results suggest similar dynamics in the lymph node.

A limitation of the estimated parameters obtained from the agent-based model is that they fit best for a specific system that describes an *in vivo* setup. As established with the PCKR and fraction of killed infected cell analyses, the results differ based on initial number of infected cells and CTLs. Thus, the parameters identified in the paper hold true for a certain experimental set-up and cannot be taken for different initial conditions. Nonetheless, the benefit of the model is the ease with which it can be adapted to various other initial conditions. Consequently, while the agent-based system described in the paper is designed for a particular *in vivo* system, it can be applied to different setting by varying the parameters and can be used to study the killing mechanisms under different conditions. Other than to study killing mechanisms, it may be used to study cell activation that is based on integrating signals from cell-cell-interactions.

## Materials and Methods

### Experimental dataset

We used the experimental dataset published by Halle et al. [14] to investigate which killing hypothesis can best explain the quantitative properties of CTL killing activity *in vivo*. Briefly, the experimental setup consists of mice infected with a modified reporter virus. We have used both murine cytomegalovirus (MCMV-3D-delta-vRAP) and modified vaccinia virus Ankara (MVA-OVA-mCherry) that both do not inhibit MHC class I peptide presentation, that both express a fluorescent protein (mCherry), and that further express a specific OVA protein-derived peptide. In parallel, we used CD8 T cells specific for the OVA-derived peptide (OT1 mouse model) that express another fluorescent protein that were adoptively transferred to the mice (prior to T cell priming and later infection with the reporter viruses). In that setting, virus-infected cells are red fluorescent while green GFP^+^ CTLs specifically recognize infected cells. Two-photon microscopy was used to observe micro-anatomical regions containing virus-infected cells inside lymph nodes. In these experiments, only a small portion of the lymph node tissue is monitored. CTLs are recruited to interact with the infected cells in this region. The movement of the T cells is tracked, together with the duration and numbers of contacts between T cells and infected cells, and finally the time at which infected cells disappear. Here, we described an agent-based model designed to reproduce this experimental setting.

### Three-Dimensional Setting and Movement of Cells

The CTLs and infected cells are positioned in a continuous three-dimensional space of 700×700×700 *μ*m(Figure 1a). The space is assumed to be periodic along x- and y-axis such that if a cell leaves from one side, it re-enters from the opposite end keeping the other coordinates same and keeping the same velocity vector. Along the z-axis the borders are impermeable, i.e. cells cannot leave from the top of the tissue. While the cells can leave through the lower boundary along z-axis in the experiments, in our simulations, we assumed a closed system along the lower and upper z-boundary. CTLs are placed randomly in the whole space whereas the infected cells are placed only in the top 40% (*Z*_lim_) of the space with respect to the z-axis, as seen *in vivo*.

Regarding modelling of CTL migration, we assumed a core repulsion of the nucleus that is assumed to be 50% of the overall size of the cell. Cells are initially positioned in the space such that no two nucliei physically overlap with each other. THe CTLs have a radius of 4.8 *μ*m (*R*_T_ and the infected cells have a radius of 5.1 *μ*m (*R*_I_) (directly taken from experimental observations).

CTLs are allowed to move and infected cells are stationary. The CTLs carry a current direction of movement and a speed value (velocity). The initial speed value is chosen from the distribution from (Figure 1d), and the initial directions are assigned randomly. The CTLs have a constant persistence time of 2 minutes which is the time that a particular cell moves in one direction before changing directions [32]. When the persistence time of a CTL is reached, it is assigned a new direction of movement and a new speed taken from the distributions obtained from experimental data. Initially, the time that each cell has been moving in the same direction is assigned as a random value between 0 and the persistence time to avoid synchronization.

### Collision Detection and Interaction between Cells

At each time step of 0.1 minutes, the following tasks are carried out:

#### (a) Collision

CTLs that are currently not interacting with an infected cell are checked for movement: If the CTL has been moving in the same direction for equal or more than the persistent time, the CTL is assigned a new speed and direction taken from the experimentally observed distribution (Figure 1d, e). The order of updating the movement of the CTLs is executed in an unsynchronized manner to avoid any bias.

Then, the CTLs are checked for collisions. The CTLs are moved in their direction of movement as far as possible within the distance they should move during that time-step, until they reach another cell, according to nucleus hard-core repulsion. More precisely, a cell is reached when the distance between the center of the CTL being moved and the other cell is *C*_CF_(*R*_T_ + *R*_I_) in the case of when the other cell is an infected cell and is *C*_CF_(*R*_T_ + *R*_T_) when the other ce is a CTL. Here *R*_T_ and *R*_I_ are the radii of the CTL and infected cells, respectively, and *C*_CF_ defines the fraction of the cell radius taken by the nucleus. If a collision takes places, the CTL changes its direction and resumes motion in another direction.

#### (b) Interaction initiated

The cells are checked for interaction. Each CTL can interact with just one infected cell at a given time but an infected cell can have multiple interactions simultaneously. If the distance between a free infected cell and a CTL is less than *I*_CF_(*R*_T_ + *R*_I_), where *I*_CF_ is the interaction confinement factor, interaction is initiated and the cell stops moving for the entire interaction duration. The factor for *I*_CF_ is set as 1.5 to account for the fact that cells may extend pseudopodia to interact with another cell. Note that an interaction can be initiated before cells collide into each other.

#### (c) Interaction terminated

Once an interaction has been initiated, the duration of that particular interaction is decided based on a predefined distribution. The experimental data for duration of interaction has very few points and to achieve a better sampling, the distribution of interaction duration is chosen from a fitted log normal distribution, leading to a mean of 2.1 and a standard deviation of 1.2 (Figure 1b).

When the duration of the ongoing interaction between a CTL and an infected cell exceeds the interaction time assigned in the beginning, the interaction is broken off and the CTL is assigned a new speed and turning angle. Further interactions are not allowed for the CTL during the first persistence time.

#### (d) Hypotheses for target cell death

Infected cells that are interacting with CTLs are checked for death. Target cells for which the cell death decision has been initiated do not vanish from the system immediately but persist for a certain time to die called *T*_death_. The value of *T*_death_ is treated as an unknown parameter and is taken to be a constant value. The interaction is terminated when either the value for *T*_death_ for the infected cell is reached or when the interaction time is completed, whichever happens first. The death decision is based on the following possible hypotheses:

1. Null hypothesis: The probability of infected cell death remains constant with increasing number of CTL contacts. The cell death decision is taken at the end of each interaction.
2. Infected cell contact integration: The probability of target cell death increases linearly with increasing number of CTL contacts. At each contact, the probability of cell death is given by *k*_I_ * *C*_I_ where *k*_I_ is constant and *C*_I_ is the number of contacts that the infected cell has had including the current one. Similar to the Null hypothesis, the cell death decision is taken once at the end of each interaction.
3. CTL contact integration: the killing capacity of the T cell is modulated with increasing number of contacts. For the case of positive modulation, at each contact, the probability of cell death is given by *k*_I_ * *C*_I_ where *k*_T_ is constant and *C*_T_ is the number of contacts that the CTL has had. Here, the CTL gets more lethal with increasing number of contacts with infected cells. On the other hand, a negative modulation of CTL killing capacity is given by *k*_T_/*C*_T_ where an increasing number of contacts with infected cells leads to a long-term exhaustion of the T cells. Death decision is taken at the end of each interaction.
4. Damage: As a possible detailed mechanism by which infected cells get more susceptible following CTL attack, infected cells accumulate damage. This damage is assumed to be proportional to the duration of contact and is computed at each time step during every interaction for an infected cell according to equation (1):

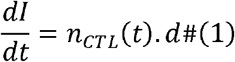

where *I* is the damage of infected cell, *d* is the damage rate and *n*_CTL_(*t*) is number of CTLs that are interacting with the infected cell at time *t*. From equation (1), it can be seen that for an infected cell to reach a damage of 1, a total contact with CTLs of 1/*d* minutes is required. For an infected cell, this is the minimum duration of contact after which it initiates apoptosis. Once an infected cell reaches a damage of 1, the death process is initiated.
5. Saturated damage: Damage of infected cells during a single interaction with CTLs is limited to a certain duration *T*_max_. For longer interactions, the CTLs cease to damage the infected cells. For each CTL, the damage induced in a particular infected cell during interaction is given by:

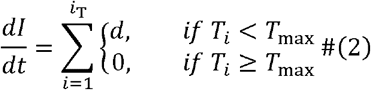

where *I* is the damage of the infected cell, *d* is the damage rate, *T*_i_ is the time duration of the *i*^th^ interaction and *i*_T_ is the total number of CTL contacts that the infected cell is currently having. Similar to the damage hypothesis outlined above, an infected cell needs an interaction of at least 1/*d* minutes to acquire a damage of 1. But if the value of 1/*d* exceeds *T*_max_, the infected cell will accumulate the damage over a series of contacts. Thus, the time to reach a damage of 1 would also include the time between two consecutive contacts and would be greater than 1/*d*.
6. Damage and repair: In addition to damage by CTLs, infected cells can also repair themselves. The repair rate is constant. At each time step time, the damage for an infected cell is updated according to:

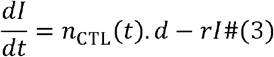

where *I* is the current damage of the infected cell, *d* is the damage rate, *n*_CTL_(*t*) is the number of CTLs interacting with the infected cell at time *t* and *r* is the repair rate.

The damage evolution during the first contact follows equation (4):

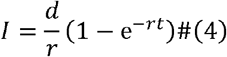

Then, the duration of the first contact to reach a damage of 1 (*t*_complete_) is given by:

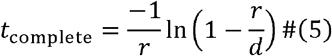

Contacts between CTLs and already dying cells, i.e. a target cells during the *T*_death_, are named “zombie contacts”. In the simulations, zombie contacts can be allowed or not, in which case CTLs ignore the dying cells as soon as the cell death decision has been established for that target cell. If zombie contacts are allowed, they do not contribute to the cell death decision but are nonetheless observed as contacts when analysing the data.

After *T*_death_ death elapses, the dying cell vanishes from the simulation.

### Finding best parameter sets

To compare the model results with the experimental results, we compared the following 8 datasets (see Figure S1):

1. Observed probability of killing infected cells with a particular number of interactions with CTLs (Figure S1a). In the model, the number of infected cells that died at exactly the *i*^th^ interaction is normalized by the total number of infected cells with at least *i* interactions.
2. Distribution of the observed time between first CTL contact and the observed time of cell disruption (*T*_elimination_) (Figure S1b). In the model, this distribution is obtained by monitoring all cells from their first contact to a CTL, counting the number of killed cells in time bins, and normalising these with the total number of killed cells.
3. Distribution of the number of contacts with CTLs for infected cells that died during the observation period (Figure S1c). In the model, the number of CTL contacts is saved with each cell and a histogram is generated at the end of the simulation and normalized with the total number of dead cells at the end of the simulation.
4. Distribution of the number of contacts with CTLs for infected cells that survived the observation period (Figure S1c). In the model, same procedure as in point 3 is used but for cells alive at the end of the simulation.
5. Total duration of contact with CTLs for infected cells that died during the observation period (Figure S1d). In the model, each cell in contact to CTL increases a clock and a histogram on time bins is generated at the end of the simulation and normalized by the total number of dead cells at the end of the simulation.
6. Total duration of contact with CTLs for infected cells that survived the observation period (Figure S1d). In the model, same procedure as in point 5 but for cells alive at the end of the simulation.
7. Distribution of single CTL contact durations for infected cells that died during the observation period (Figure S1e). In the model, at each contact with CTLs a clock is running and the time when the cells detach is saved. For all dead cells, these times are recollected in a histogram on time bins at the end of the simulation and normalized with the total number of contacts of all dead cells.
8. Distribution of single CTL contacts duration for infected cells that survived the observation period (Figure S1e). In the model, same procedure as in point 7 but for cells alive at the end of the simulation.

Although most parameters are directly taken from data, each hypothesis carries its set of unknown parameters, like a killing probability or damage rate, and *T*_death_. For each parameter set, we ran 30 simulations and calculated the respective cost for each of the 8 datasets described above and for each dataset the average cost was calculated over these 30 simulations. This gave us 8 values which were the average costs for each of the 8 datasets described above. When comparing different parameter sets, the formula in equation (6) is used to compute the cost for each of the 8 datasets described above:

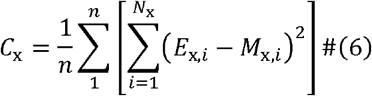

where *C*_x_ is the average cost of the *x*^th^ dataset, *n* is the total number of simulations run for each parameter set, *N*_x_ is the total number of data points of the *x*^th^ experimental dataset, *E*_x,i_ is the *i*^th^ data point of the *x*^th^ experimental dataset and *M*_x,*i*_ is the *i*^th^ data point of the *i*^th^ model dataset with this parameter set. The subscript *x* represents the 8 datasets described above.

The 8 separate costs were averaged to compute the mean cost for a particular parameter set using equation:

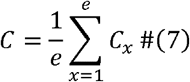

where *C* is the average cost for a particular simulation set, is the total number of datasets (*e* = 8 in the current study).

For each hypothesis, the average costs are plotted in a heatmap comparing different combination of parameter values. Then, the parameter set with the lowest cost for each hypothesis was used to run the simulation to get the results. For the first dataset, since the experimental data points beyond the 6^th^ contact only represent a small number of cells, only the 6 first interactions are included in the cost calculation.

Table of Parameters Fixed from Data

**Table of Parameters Fixed from Data**
| Parameter | Value |
| --- | --- |
| Number of CTLs ($N_T$) | 200 |
| Number of infected cells ($N_I$) | 250 |
| Percent of z-axis occupied by infected cells ($Z_{lim}$) | 40% |
| Radius of CTLs ($R_T$) | 4.8 $\mu\text{m}$ |
| Radius of infected cells ($R_I$) | 5.1 $\mu\text{m}$ |
| Interaction confinement factor ($I_{CF}$) | 1.5 |
| Collision confinement factor ($C_{CF}$) | 0.5 |
| Persistent time ($T_{pers}$) | 2 minutes |
| X-axis dimension of space ($X_{dim}$) | 700 $\mu\text{m}$ |
| Y-axis dimension of space ($Y_{dim}$) | 700 $\mu\text{m}$ |
| Z-axis dimension of space ($Z_{dim}$) | 700 $\mu\text{m}$ |
| Simulation ($T_{Sim}$) | 240 minutes |
| Speed | Taken from distribution (Figure 1d) |
| Turning angle | Taken from distribution (Figure 1e) |
| Interaction duration | Taken from distribution (Figure 1f) |

## Supplement

**Figure S1:**
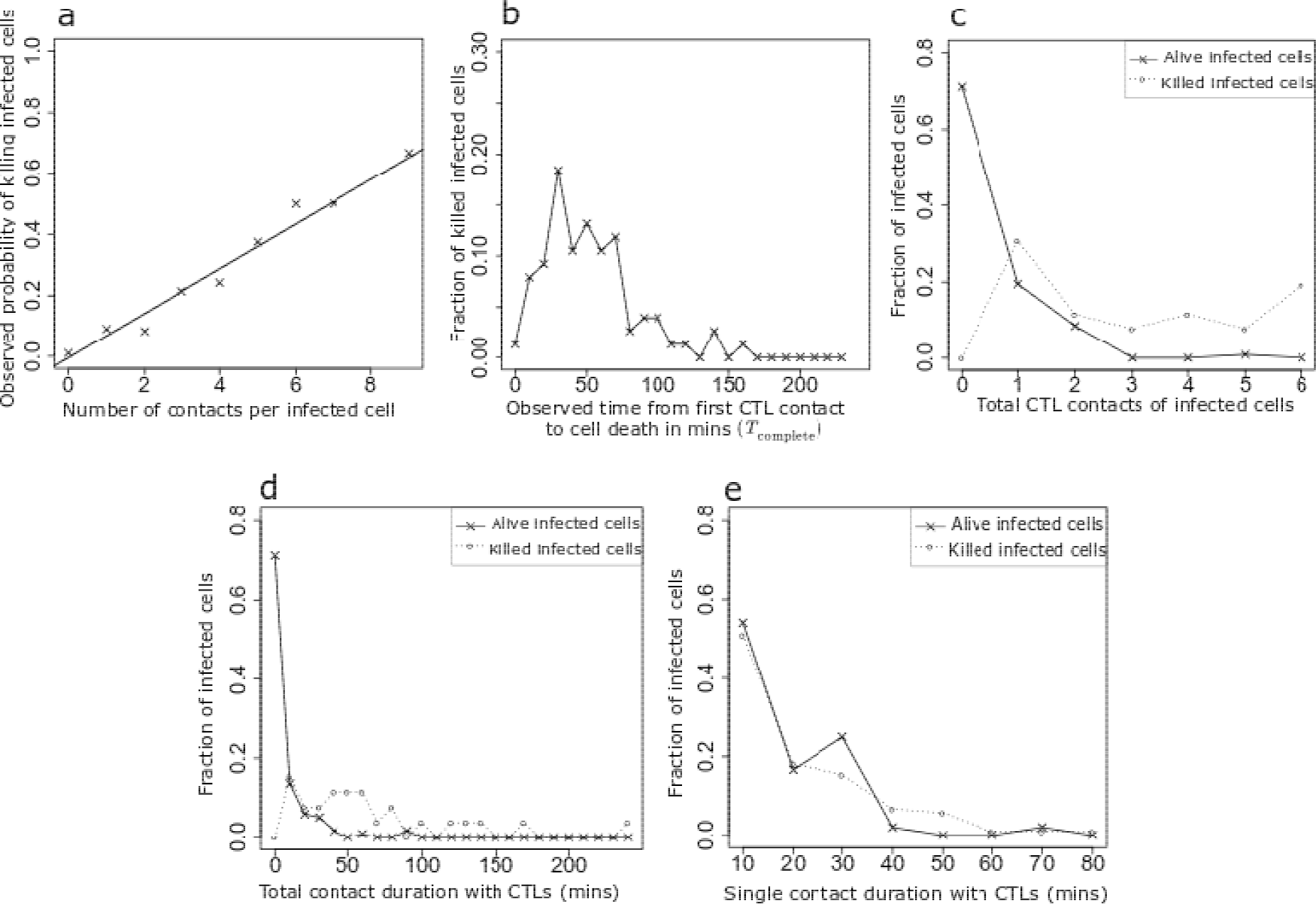
Experimental readouts used to compare experimental data with simulation data. (a) Fraction of killed infected cells at each interaction, (b) Distribution of observed times between the first contact to a CTL and the actual cell death for all infected cells that were killed, (c) Distribution of the number of contacts with CTLs for all killed and alive infected cells during the observation period. (d, e) Distribution of total (d) and single (e) contact durations with CTLs for killed and alive infected cells during the observation period.

**Figure S2:**
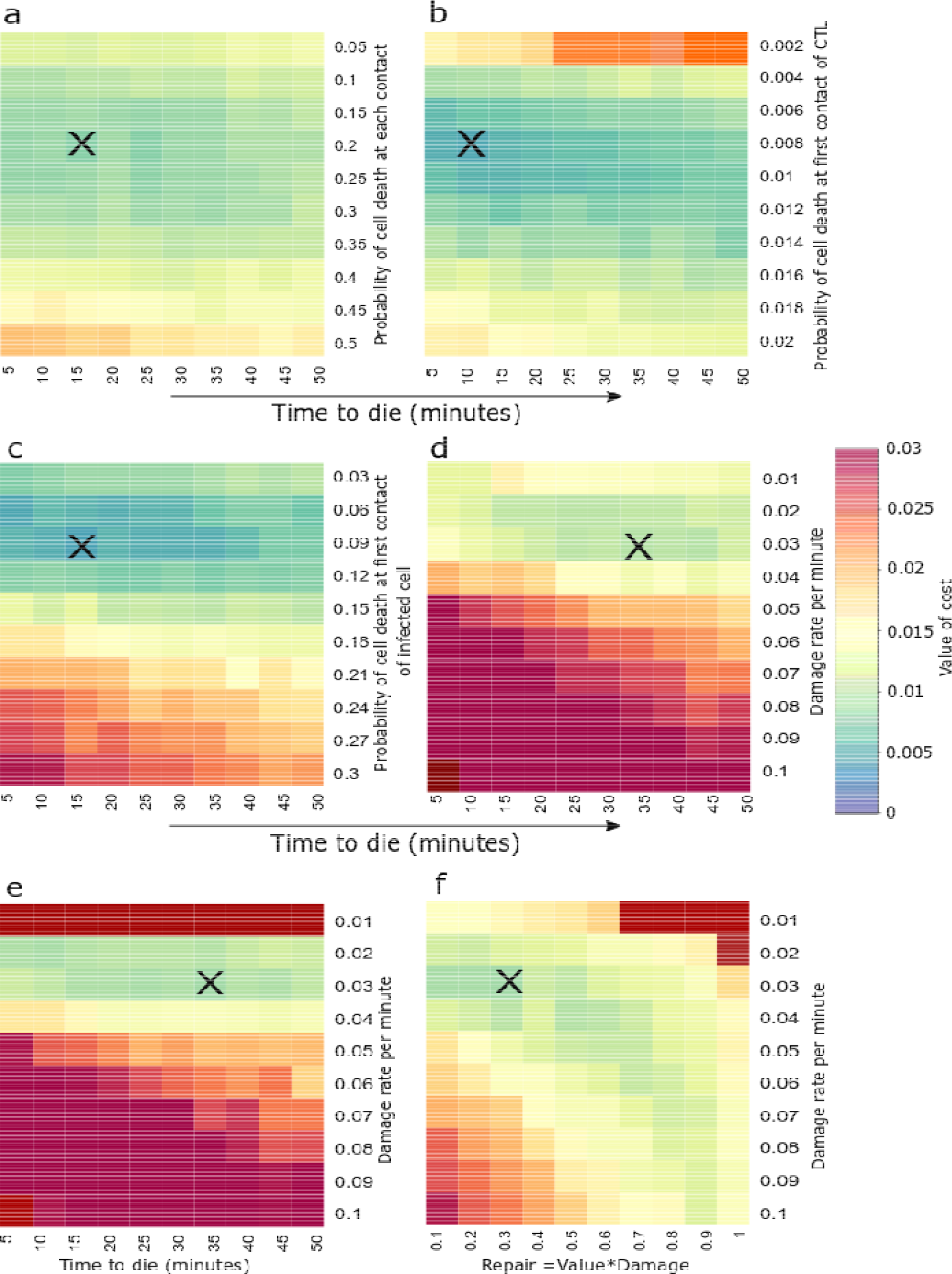
Heatmaps for all hypotheses in absence of zombie contacts. (a) Null hypothesis; (b) CTL contact integration hypothesis; (c) Infected cell contact integration hypothesis; (d) Constant damage hypothesis; (e) Saturated damage hypothesis; (f) Concomitant damage and repair hypothesis. Each point on the heatmap is obtained by calculating the average cost over 30 simulations for the respective parameter combination. ‘X’ represents the parameter combination with the lowest cost. For saturated damage hypothesis and damage and repair hypothesis, there are three variable parameters and the lowest costs were found scanning the 3D parameter space (data not shown).

**Figure S3:**
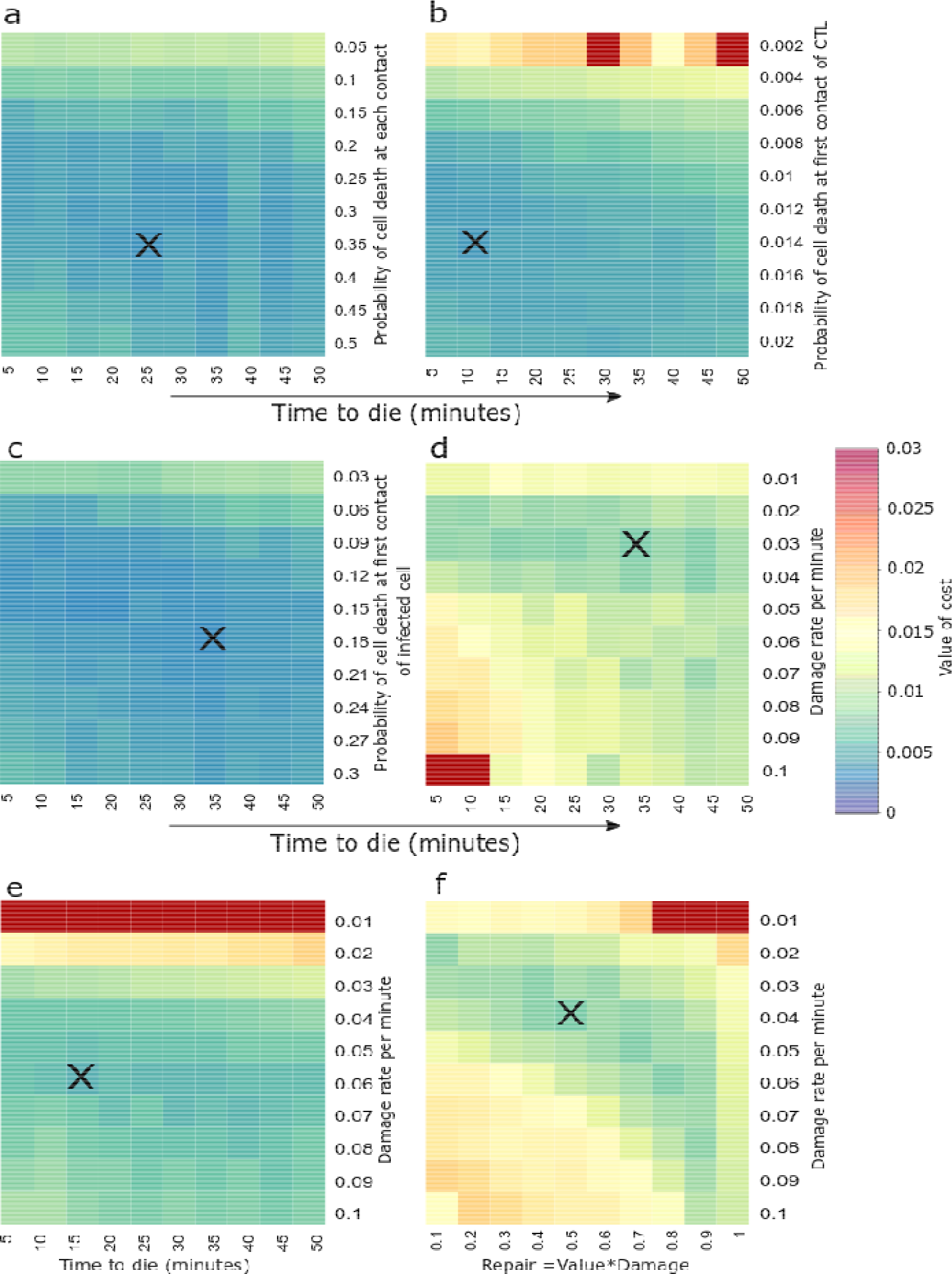
Heatmaps for all hypotheses in presence of zombie contacts. (a) Null hypothesis; (b) CTL contact integration hypothesis; (c) Infected cell contact integration hypothesis; (d) Constant damage hypothesis; (e) Saturated damage hypothesis; (f) Concomitant damage and repair hypothesis. The heatmaps are obtained using the same conditions described in Figure S2.

